# Large-scale nuclear remodeling and transcriptional deregulation occur on both derivative chromosomes after Mantle Cell Lymphoma chromosomal translocation

**DOI:** 10.1101/2019.12.30.882407

**Authors:** Fatimata Bintou Sall, Andrei Pichugin, Olga Iarovaia, Ana Barat, Tatyana Tsfasman, Caroline Brossas, Marie-Noëlle Prioleau, Eugeny V. Sheval, Anastasiya A. Zharikova, Julien Lazarovici, Valérie Camara-Clayette, Vincent Ribrag, Marc Lipinski, Yegor Vassetzky, Diego Germini

**Affiliations:** UMR9018, Université Paris-Saclay, CNRS, Gustave Roussy, Villejuif, France; Laboratoire d’Hématologie, Centre Hospitalier Universitaire Aristide Le Dantec, Université Cheikh Anta Diop, Dakar, Senegal; Institute of Gene Biology, Russian Academy of Sciences, Moscow, Russia; UMR7592, Université Paris Diderot, CNRS, Institut Jacques Monod, Paris, France; Belozersky Institute of Physico-Chemical Biology, Moscow State University, Russia; Département d’Hématologie, Gustave Roussy, Université Paris-Saclay, Villejuif, France; AMMICa INSERMUS23/CNRS UMR3655, Translational Hematology Unit, Gustave Roussy, Villejuif, France; Département des Innovations Thérapeutiques et Essais Précoces, Gustave Roussy, Université Paris-Saclay, Villejuif, France; Koltzov Institute of Developmental Biology, Russian Academy of Sciences, Moscow, Russia

## Abstract

Recurrent chromosomal translocations are found in many blood and solid cancers. Balanced translocations, frequent in lymphoid malignancies, lead to the formation of two aberrant derivative (der) chromosomes. This event often leads to overexpression of an oncogene. In many cases, the expression of an oncogene is not enough to produce a malignant phenotype; however, most part of the studies focus on the events involving the chromosome where the oncogene is located, but rarely the other der chromosome where other oncogenic alterations may potentially arise. Mantle cell lymphoma (MCL), an aggressive B-cell non-Hodgkin lymphoma, is a perfect example of this. In 85% of the cases, it is characterized by the translocation t(11;14), which leads to the overexpression of cyclin D1 (*CCND1*) gene which results juxtaposed to the immunoglobulin heavy chain (*IGH*) gene on the der14 chromosome. This feature alone is not sufficient to induce oncogenesis. Here we focused on the der11 chromosome. We demonstrated that expression of 88 genes located in a 15mb region close to the translocation breakpoint on the der11 was deregulated both in the GRANTA-519 MCL cell line and in B-cells from MCL patients. We found that a large segment of der11containing deregulated genes was relocated from its normal position in the nuclear periphery towards the center of the nucleus in close proximity to the nucleolus where the abundant nucleolar protein nucleolin binds a subset of genes located close to the breakpoint and activates their expression. This finding allowed to identify new potential oncogenes involved in MCL and the mechanisms of their upregulation.

## INTRODUCTION

Mantle cell lymphoma (MCL) is a non-Hodgkin lymphoma characterized by an uncontrolled proliferation of the B-cells located in the outer edge of a lymph node (mantle zone). It represents 8% of all lymphomas in Europe (Vose, 2017). It has a poor prognosis with an average survival rate of five years (Skarbnik and Goy, 2015). Despite significant advances in treatment (Camara-Clayette et al., 2012), MCL still remains an incurable disease (Campo and Rule, 2015). Moreover, its annual incidence has increased during recent decades (Epperla et al., 2017).

It is characterized by a specific t(11;14)(q13; q32) (Bentz et al., 2000; Li et al., 1999) translocation where the 11q13 region (encoding for over 300 genes) is recombined with the Immunoglobulin Heavy Chain (*IGH*) locus on chromosome 14.

This translocation leads to the over-expression of the gene encoding for the G1/S specific Cyclin D1 (*CCND1*) that is not expressed in quiescent B lymphocytes (Bosch et al., 1994; Dreyling et al., 1997).

However t(11;14) can be also detected in healthy people’s blood cells (Hirt et al., 2004) and expression of *CCND1* under the control of different known *IGH* enhancers in transgenic mice is not sufficient for tumor development (Fiancette et al., 2010; Lovec et al., 1994). Indeed it seems that *CCND1* is not the only gene affected in MCL and recurrent mutations are found in other genes belonging to different pathways and cellular processes e.g. cell cycle (*ATM*), epigenetic regulation (*MLL2* and *3*), NF-kB pathway (*BIRC2* and *3*) (reviewed in (Ahmed et al., 2016)). Some MCL cases exist lacking *CCND1* overexpression but carrying secondary alteration as *SOX11*overexpression (Jares et al., 2012). These data point out that additional oncogenic alterations are necessary to develop a malignant phenotype.

Data previously obtained in our laboratory show that the *CCND1* locus, normally located at the periphery of the nucleus, after the t(11;14), in MCL cell lines or in lymphoma cells from patients, is regularly found close to the center of the nucleus and/or associated to the nucleolus. In the surroundings of the nucleolus, *foci* of active polymerase II, therefore regions of transcriptionally active chromatin, and a range of potential transcription factors have been found. For instance nucleolin, a major nucleolar protein, which forms, together with hnRNP D, the B cells-specific heterodimeric transcription factor LR1, may be involved in activation of oncogenes in lymphomas (Allinne et al., 2014).

Following a chromosomal translocation, changes in spatial gene localization and nuclear reorganization occur, inducing alterations in gene expression and potentially leading to additional events necessary for oncogenesis (reviewed in (Harewood and Fraser, 2014)). In the present work, we looked for possible oncogenic modifications in genomic regions other than the *CCND1* and the der14 chromosome and we focused our attention on the events involving the der11 chromosome where only a small portion of the chromosome 14 is translocated.

We describe here new features specific of MCL and potentially linked to the malignant transformation. We have shown in MCL cells (*in vitro* and from MCL patients) a massive deregulation of the expression of genes located on the chromosome 11 close to the translocation breakpoint. Most of these genes were located on the der11 after the t(11;14). We explained this event by showing that the der11 chromosome is relocated in close proximity to the nucleolus where the nucleolin directly binds to chromatin, modulating gene expression. The same nuclear reorganization was also found in B-cells from MCL patients harboring the t(11;14). Our results describe novel genomic and transcriptomic alterations for MCL, a possible role played by nucleolin in inducing potential lymphomagenic alterations and provide potential new targets for the amelioration of the prognosis for MCL.

## RESULTS

### Analysis of clustered gene expression in MCL reveals a large domain of upregulated genes adjacent to the translocation point on der11chromosome

We first wanted to determine whether the t(11;14) perturbs the expression of genes other than *CCND1* in the vicinity of the translocation breakpoint.

We have performed RNA-seq of MCL cell lines (GRANTA-519, carrying the t(11;14)) and RPMI8866 lymphoblastoid cell lines (LCL) used as a control. Gene expression in GRANTA-519 was compared to LCL RPMI8866 and the genes significantly upregulated (adjusted p value<0.05) were mapped on each chromosome. We found that a total of 286 genes were significantly upregulated on the chromosome 11. After the translocation, the portion of genes located on der11 in close proximity to the *CCND1* (Figure 1A, in green) shows a more deregulated pattern as compared to the genes located on the der14 (Figure 1A, in red). Dividing the chromosome 11 in equal portions of 15mb, we found that the greatest percentage of upregulated genes was located in the region surrounding the translocation breakpoint compared to the rest of the chromosome (Figure 1B). That region contains genes mainly located on the der11. Such a relatively high transcriptional deregulation region cannot be found within other chromosomes (**Supplementary Figure 1**).

**Figure 1.**
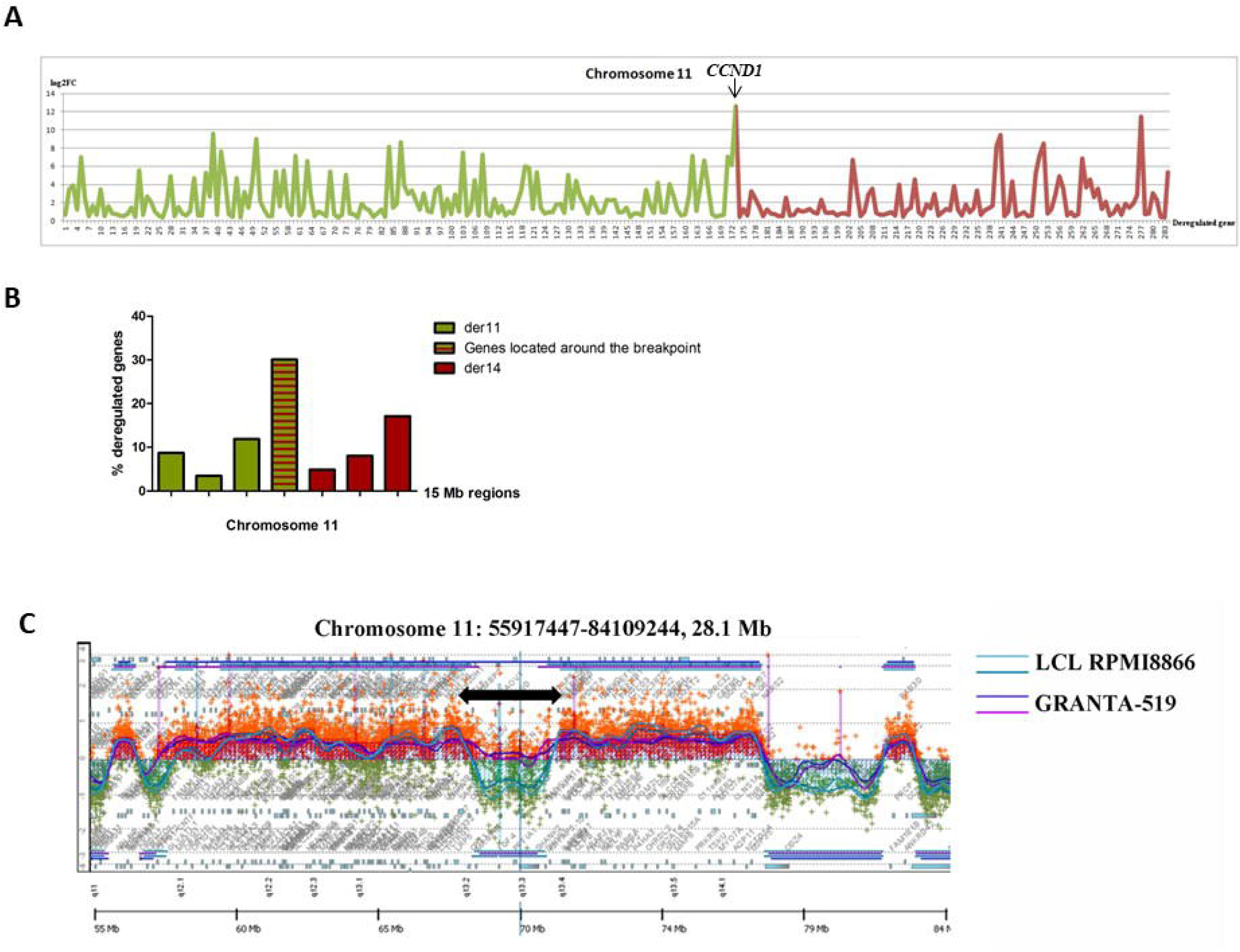
Gene transcription and replication timing is perturbed on the chromosome 11 after the t(11;14) translocation. **(A)** Profile of gene transcription upregulation along chromosome 11 in GRANTA-519 cells compared to LCL RPMI8866. RNA was extracted from GRANTA-519 cells and LCL RPMI 8866, used as a control, and RNA seq performed. Upregulated genes were mapped along the chromosome 11, on the Y axis log2 fold change (FC) of each gene is reported from the first upregulated gene (“1” on the X axis) on the p arm of the chromosome 11 to the last one on the telomeric region. Log2FC of the genes located upstream (on the der11 after the t(11;14)) and downstream (on the der14 after the t(11;14)) of the *CCND1* are represented in green and red, respectively. **(B)** Percentage of deregulated genes in seven non overlapping 15mb regions of the chromosome 11. The percentage was calculated on the total number of upregulated genes of the chromosome 11. Genes located upstream (on the der11 after the t(11;14)) and downstream (on the der14 after the t(11;14)) of the *CCND1* are represented in green and red, respectively. The red and green histograms represent the percentage of upregulated genes in the 15mb surrounding the *CCND1*. **(C)** Replication timing profiles at the region surrounding the translocation breakpoint on the chromosome 11. The profiles are represented with two light blue lines for LCL RPMI8866 and two violet lines for MCL cells GRANTA-519. Cells were sorted into two fractions of S-Phase (early and late S-Phase). Pulse labeled BrdU nascent DNA from each fraction was recovered by immunoprecipitation and differentially labeled before co-hybridization to a human genome CGH microarray, containing one 60-mer oligonucleotide probe every 13 Kb. The log2-ratio (early/late) of the abundance of each probe in the early and late S-phase is shown. The timing profiles were smoothed using the Moving Average option of the Agilent Genomic Workbench 6.5 software with the triangular algorithm and 500 kb windows. Fast replicating genes are represented in red while in green the slow replicating ones. The black arrow highlight region where the replication timing shift is more evident.

Replication timing of genomic regions correlates with gene transcription and changes in nuclear position. Active transcription coincides with a switch from late to early replication timing (Rhind and Gilbert, 2013; Rivera-Mulia et al., 2015). In order to verify this correlation in our experimental model, we analyzed the DNA replication timing in GRANTA-519 and in LCL RPMI8866 as a control. Cells labeled with BrdU were sorted into two S-Phase fractions (early and late S-Phase) and immunoprecipitated with an anti-BrdU antibody, amplified, differentially labeled, and co-hybridized onto a whole human genome oligonucleotide microarray. The replication-timing profile represented here is obtained by the log_2_-ratio of the abundance of each genomic probe in the early and late S-phase fractions, revealing early and late replicated domains.

We observed a significant shift to early replication timing surrounding the *CCND*1 gene in GRANTA-519 (purple lines) in contrast to the LCL RPMI8866 (blue lines) (Figure 1C). This 10 Mb modified region (black arrow in Figure 1C) moves from a late S-Phase to a Mid-S-Phase replication timing (Figure 1C).

The replication timing modification covers a wide region upstream and downstream the *CCND*1 gene covering from the 11q13.2 to the 11q13.4 loci. This region coincides to the one where the massive gene deregulation was observed in the RNA-seq, thus supporting our data. The replication timing profile at the *CCND*1 locus corresponds to an average of the replication timing of the two alleles (wild type and translocated), leading to an underestimation of the replication timing shift due to the translocation. For this reason, we can assume that the replication timing of the translocated allele of chromosome 11 at the translocation breakpoint moves from late-S-phase to early-S-phase, while the other allele keeps the same late replication timing as the LCL RPMI8866.

To understand if the data obtained *in vitro* has relevance *in vivo*, we performed RNA-seq on B-cells purified from four MCL patients in leukemic phase. Eighty-eight genes were commonly upregulated in GRANTA-519 and in the four patients compared to the LCLRPMI8866 (Figure 2). Forty-eight percent of them (42 genes) were located in the same 15mb region surrounding the translocation breakpoint while the remaining 52% was distributed throughout the whole chromosome 11 (Figure 2).

**Figure 2.**
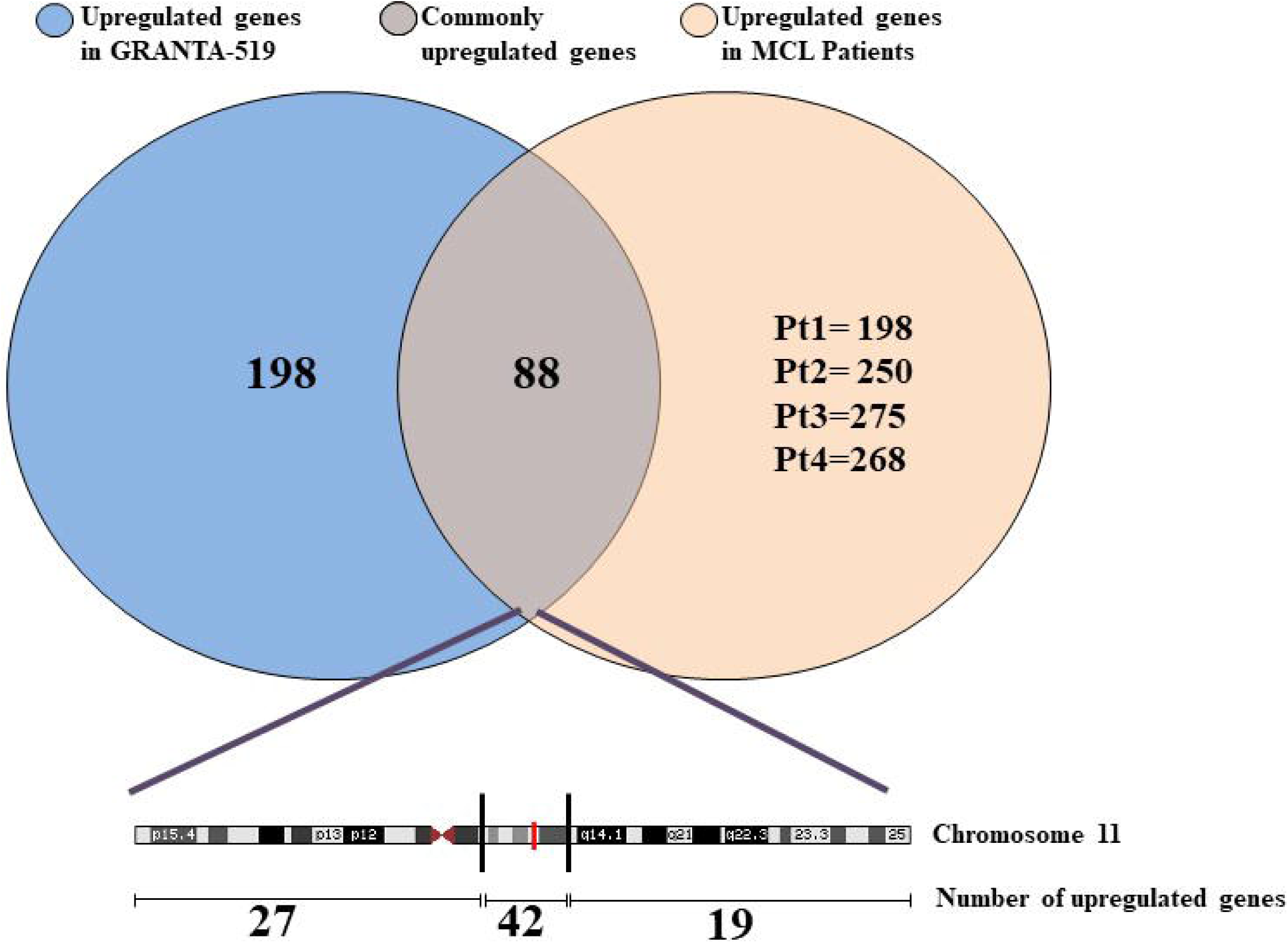
RNA-seq in GRNATA-519 cells and in B-cells from MCL patients reveals commonly upregulated genes. RNA-seq was performed on RNA extracted from GRANTA-519 cells, B-cells purified from MCL patients and LCL RPMI8866. All the transcriptomes were compared with the transcriptome of LCL RPMI8866. Upregulated genes from all comparisons were mapped on the different chromosomes. Upregulated genes located on the chromosome 11 are represented. The overlapping area among GRANTA-519 and the four MCL patients represents the commonly upregulated gene (n=88). At the bottom is schematically represented where these genes are distributed along the chromosome 11. The two black lines delimit the 15mb region surrounding the translocation breakpoint (represented by a red line). All RNA-seq were performed in triplicates.

Among the deregulated genes located on the der11 close to the breakpoint there are several that has been correlated to multiple cancer development such *MEN1* implied in chromatin remodelling, *RAD9A* involved in DNA damage response, *CTSF* involved in apoptosis regulation (Bassi et al., 2016; Covington and Fuqua, 2014; Ji et al., 2017; Lee et al., 2004),*GSTP1* found deregulated in other cases of MCL and implicated in the cell detoxification from xenobiotics (Yuan et al., 2008).

This global overexpression pattern specific of MCL describes transcriptional alterations other than *CCND1* and can hardly be explained by the action of a single enhancer but rather could be a result of a large-scale post-translocation modification. Alterations in the nuclear position of chromosomal loci can explain such transcriptional modifications due to a newly acquired vicinity to nuclear region rich in transcriptional factor or transcriptionally active as the nucleolus. Thus, we next investigated the nuclear positioning of translocated and non-translocated loci.

### Der11 chromosome relocates next to the nucleolus after the t(11;14)

We have studied, using the immuno-3D-FISH technique, the nuclear organization in MCL (GRANTA-519) and control cell lines LCL RPMI8866 and its changes after the t(11;14). Der11 chromosome was visualized in 3D-FISH as colocalized signals of *GSTP1* gene locus (colored in green) located on chromosome 11 upstream to the translocation breakpoint and the telomeric portion of the chromosome 14 (“Tel14*IGH*” - in red) (**Supplementary Figure 2**). We divided the nucleus in 10 concentric volumes where the volume “1” represents the periphery of the nucleus and the volume "10" represents its center. As shown in Figure 3A (top panels) in control cells LCL RPMI8866, in the absence of t(11;14), the *GSTP1*locusis mostly found between the center and the periphery of the nucleus (being mostly located in volume "5") while the *IGH* locus is closer to the nucleus center (mostly located in volumes "7, 8"). In GRANTA-519 cells the same loci located on the der11 chromosome relocate as compared to control cells, both moving towards the nucleus center (mostly located in volumes "8-9", Figure 3A bottom panels).

**Figure 3.**
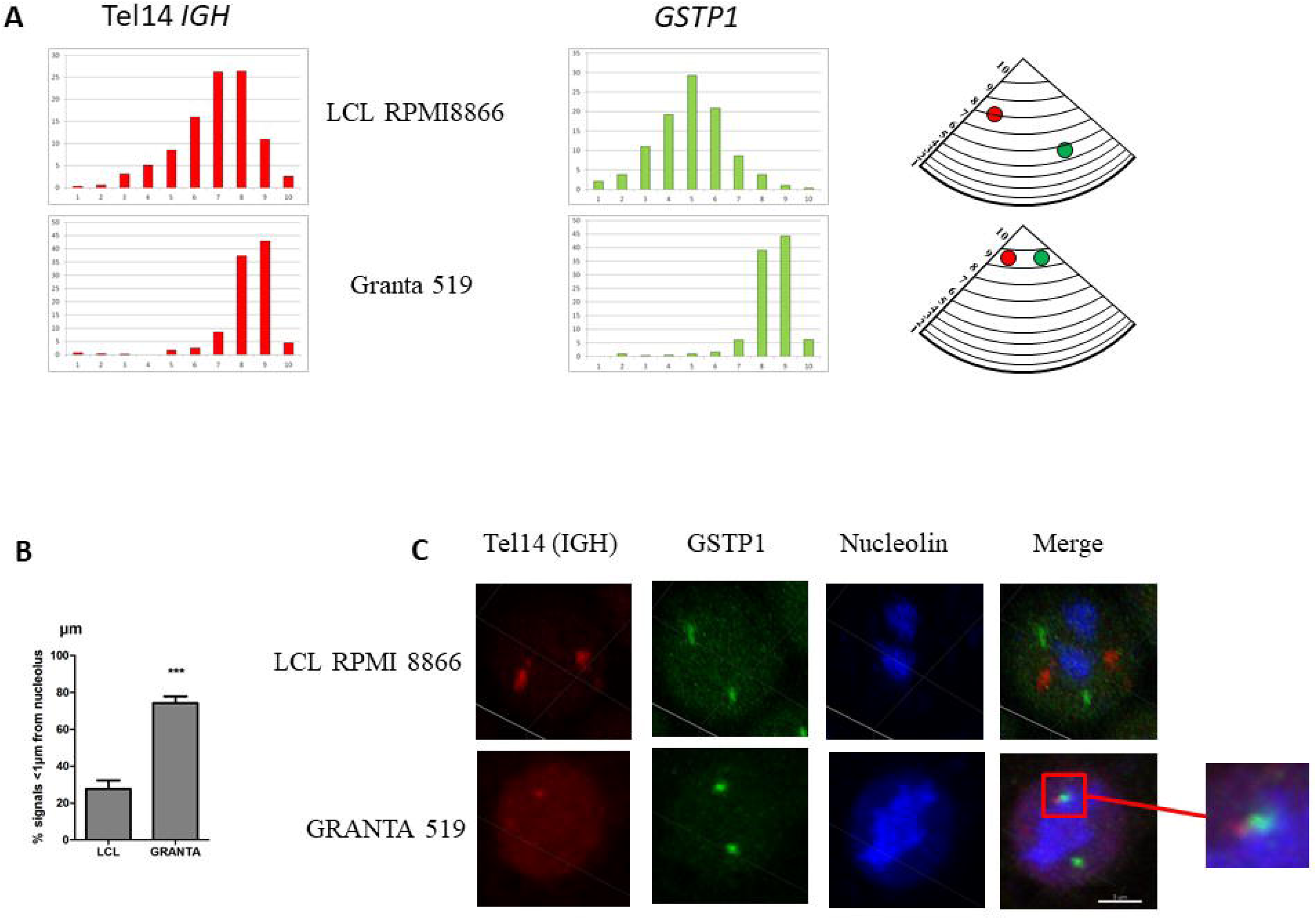
After the t(11;14), translocated alleles move in the nuclear space in close proximity with the nucleolus. (**A**) Radial distribution of the terminal part of the *IGH* gene locus (chromosome 14) and of the *GSTP1* locus (chromosome 11) in nuclei from MCL cells GRANTA-519 (where at least one allele of these loci is translocated in the t(11;14)) and LCL RPMI8866 where they do not translocate. X axis, distribution of fluorescent foci in 10 nuclear fractions of equal volume from the periphery ("1", left) to the center ("10", right) of the nucleus; Y-axis, percentage of fluorescent foci in corresponding fraction. The radial position of *IGH* and *GSTP1* loci is schematically represented in the two pictures on the right. The triangle represents a portion of the nucleus divided into 10 equal volumes. The red (*IGH*) and green (*GSTP1*) dots are placed in the nuclear fractions where the highest amount of signals was found. (**B**) Percentage of signals whose distance to the nucleolus is lower or equal to 1µm. Results are represented as means ± SEM. ***: p<0.001. A minimum of 216 loci were analyzed. (**C**) Representative 3D-FISH images showing non translocated (LCL RPMI8866, top panels) and colocalized/translocated (GRANTA-519, bottom panels) *IGH* (red) and *GSTP1* (green) gene loci in cell nuclei. Red and green colocalized signals determined the translocated alleles (der11). The square in the lower merge panel points to colocalized alleles. Nucleolus is visualized by nucleolin staining and represented in blue. Scale bar, 5 µm.

We have previously shown that in MCL and other cancer cells, the perinucleolar region is enriched in active transcription zones called «transcription factories» (Allinne et al., 2014; Razin et al., 2011). The movement towards transcriptional active areas could correlate with transcriptional deregulation as that observed in genes located on the der11. Thus, we checked if the *GSTP1* movement in the nuclear space after the translocation corresponds to a relocalization towards the nucleolus. We measured and compared the distance of each *GSTP1* signal colocalized, in GRANTA-519 (der11), and not colocalized, in LCL RPMI8866, with Tel14*IGH*to the closer nucleolus (visualized by staining the major nucleolar protein, nucleolin). We found that in GRANTA-519 cells the percentage of cells where the *GSTP1* locus on der11 is closer than 1µm to the nucleolus is significantly higher than in control LCL RPMI8866 (74.2±3.5 *vs* 27.6±4.7) (Figure 3B). A representative 3D-FISH image of this event is shown in Figure 3C. Also the average distance from the nucleolus of the *GSTP1* loci on the der11 is 0.73±0.04 µm which is significantly closer compared to normal LCL RPMI8866 (average distance=1.94±0.07 µm) (**Supplementary Figure 3**).

The demonstrated relocalization in the nuclear space of the loci on theder11 chromosome close to the nucleolus can potentially expose them to a transcriptional activation and to the action of transcription factors that can explain the overexpression of the cluster of genes in MCL described above.

### Der11 proximity to the nucleolus is present also in circulating MCL B-cells from patients

To understand whether our *in vitro* findings could be relevant to clinical conditions, we then focused our attention on circulating B-cells isolated from MCL patients. The comparison between the position of the der11 and the non-translocated loci within the same sample was performed as described above by using the 3D-FISH technique. In three out of four MCL patients the percentage of *GSTP1* signals located at less than 1µm from the nucleolus is significantly higher for the loci on the der11 than on the intact 11 chromosome (Figure 4A). Considering the means of all the four patients, the percentage of the *GSTP1* signals closer than 1µm to the nucleolus is higher when *GSTP1* is on der11 than when is on the intact chromosome 11 (53.5±5.7% vs 44.5±4.1%, respectively) (Figure 4B).

**Figure 4.**
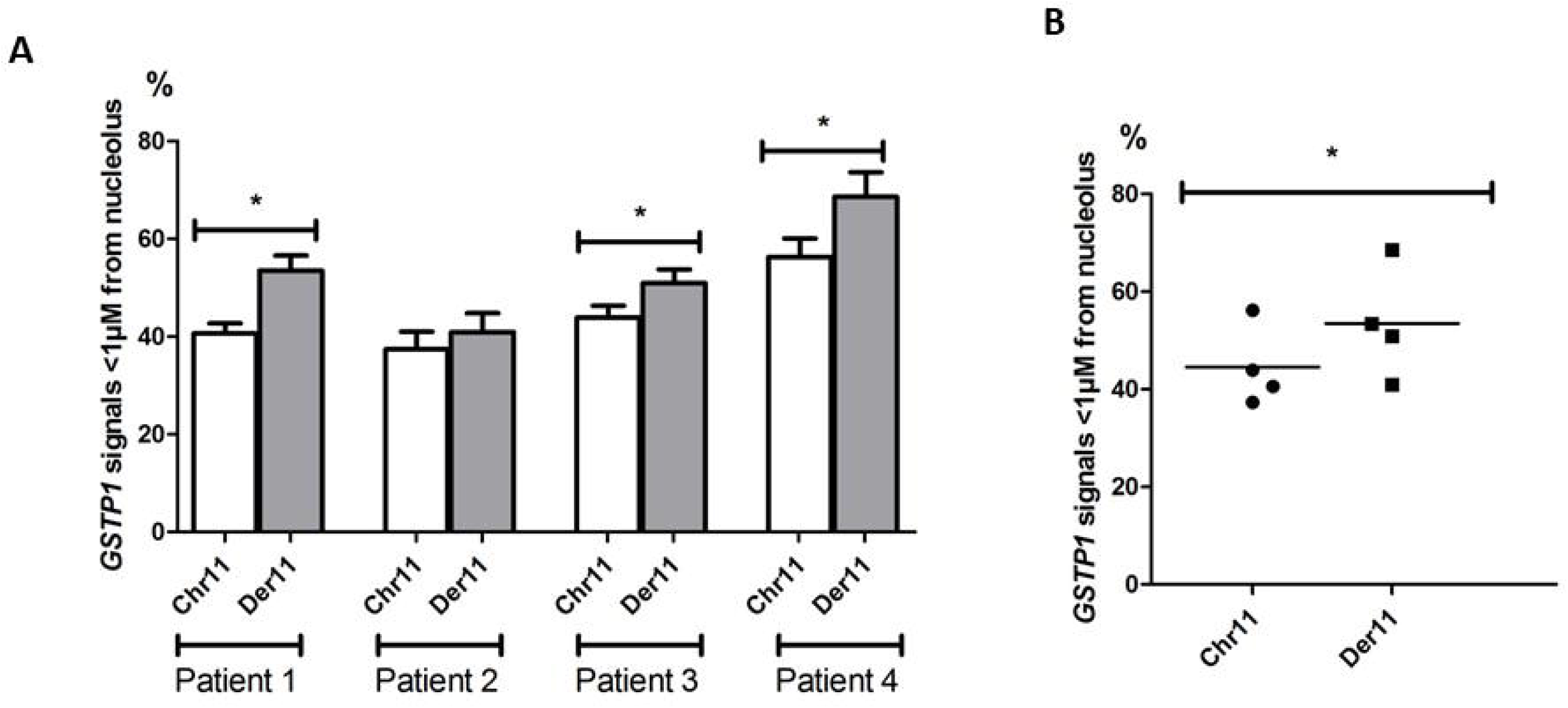
Der11 chromosome is closer to the nucleolus in MCL patients. (**A**) The percentage of translocated (Der11, grey histograms) and non-translocated *GSTP1*loci (Chr11, white histograms) closer than 1µm to nucleolus was calculated in B-cells purified from four MCL patients. *: p<0.05. (**B**) Means of these percentages were compared using the t-test. *: P<0.001.

These results obtained from B-cells purified from MCL patients support the previously described *in vitro* results.

### Nucleolin activates the transcription of der11 overexpressed genes

LR1 is a B-cell-specific transcriptional factor, binding a known DNA consensus sequence GNCNAG(G/C)CTG(A/G) (Hanakahi et al., 1997)and capable of regulating transcription in activated B-cells. LR1 is composed by two polypeptides of 106 and 45 kDa. The 106 kDa component is the nucleolin (Hanakahi et al., 1997).

Thus, we wanted to understand if the newly acquired proximity to the nucleolus and, consequently, to the nucleolin of the der11 chromosome could play a role on regulation of gene transcription.

We have performed ChIP-seq analysis targeting nucleolin on GRANTA-519 and LCL RPMI8866. Binding site analysis performed using MEME (Bailey and Elkan, 1994) revealed a motif consistent with the published one and a novel one (**Supplementary Figure 4**). Bioinformatic and Integrated Genome Viewer analysis revealed peaks corresponding to nucleolin binding sites in the same 15 mb region where the massive gene expression regulation was found. The number of peaks found in that region in LCL RPMI8866 and GRANTA-519 did not differ significantly (~32 on average between the two replicates). Nevertheless, peaks present only in GRANTA-519 were found in correspondence of significantly upregulated genes (Figure 5). Interestingly those genes were clusterized at the beginning (e.g. *CD6*), in the middle (e.g. *EHD1*) or close to the *CCND1* (e.g. *CPT1A*) of the 15 mb region.

**Figure 5.**
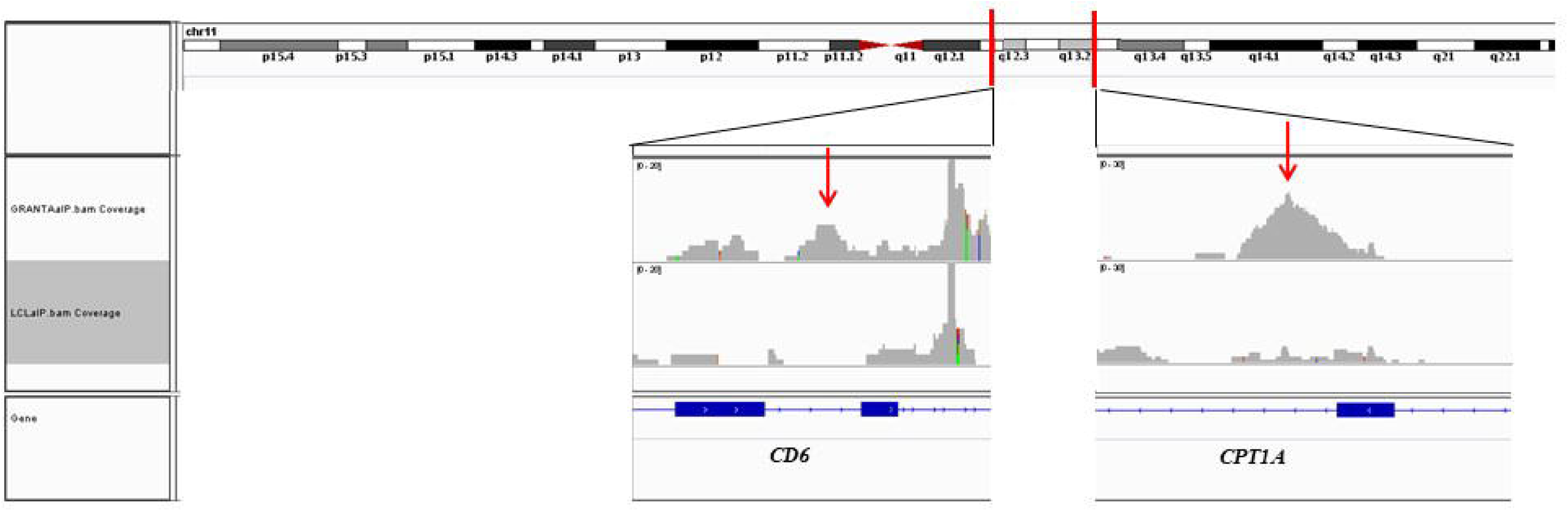
Nucleolin binds in proximity of transcriptional deregulated genes. Integrated Genome Viewer visualization of nucleolin peaks are shown in GRANTA-519(upper lane) or LCL RPMI8866 (lower lane). ChIP-seq experiment was performed in duplicate. The nucleolin profiles of the *CD6* and of the *CPT1A* locus are shown. The red lines are drawn in correspondence of the gene locus position on the chromosome 11 and the red arrows highlights the peaks observed in GRANTA-519 and not in LCL RPMI8866.

To further confirm the transactivating role of nucleolin, we ectopically expressed GFP-nucleolin in HEK cells. GFP+ cells were sorted, and the expression of genes located in the 15 mb region and commonly deregulated in GRANTA-519 cells and the four MCL patients was analyzed and compared to non-transfected HEK. In transfected cells the expression of the gene encoding for the nucleolin was 210.4±38.3 fold higher than non-transfected HEK (Figure 6A). The presence of a higher amount of nucleolin significantly increased the expression of *CD6*, *PYGM, CTSF* and*CPT1A* (Figure 6B). In correspondence of *CD6* and *CPT1A*, peaks in ChIP-seq analysis were also found. The expression of genes located on the chr11 (*DAK* and *RTN3*) or on other chromosomes (*SOD1* and *RUNX1T1*) that was unaffected in the RNA-seq remained unchanged in transfected HEK (Figure 6B, rightmost histograms), thus confirming the specific effect of the nucleolin.

**Figure 6.**
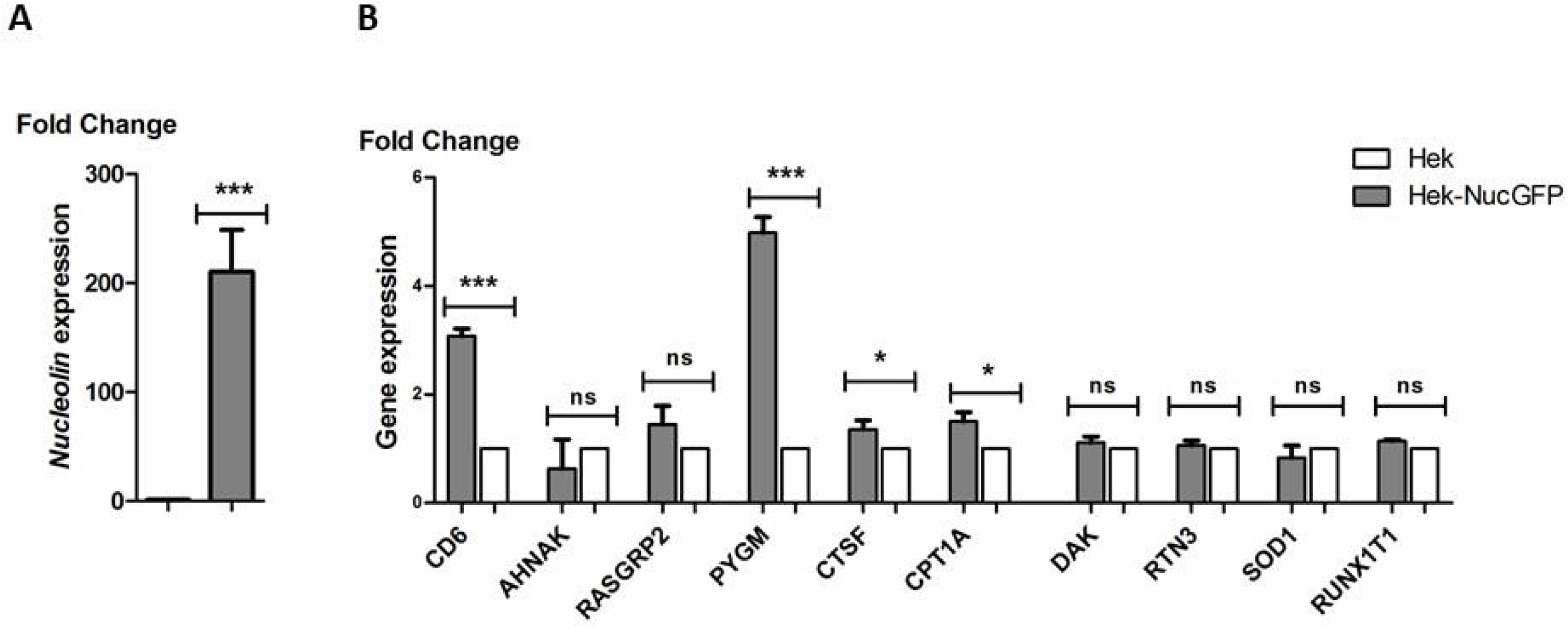
Ectopic expression of nucleolin induces overexpression of genes located close to the translocation breakpoint. Analysis of *Nucleolin* (**A**), *CD6*, *AHNAK*, *RASGRP2*, *PYGM*, *CTSF*, *CPT1A*, *DAK*, *RTN3*, *SOD1* and *RUNX1T1* (**B**) transcription in HEK cells transfected with a nucleolin encoding plasmid (grey histograms) or non-transfected (controls, white histograms). The transcription of the target genes was measured by qRT-PCR. Fold change expressions are in comparison with controls and after normalization vs. *GAPDH*. *CD6*, *AHNAK*, *RASGRP2*, *PYGM*, *CTSF*, *CPT1A* are located on the chromosome 11 and upregulated in the RNA-seq analysis, *DAK*, *RTN3,SOD1* and *RUNX1T1* served as controls as they were not deregulated in the RNA-seq analysis and are located on chromosomes 11 (*DAK*, *RTN3*), 21 (*SOD1*) and 8 (*RUNX1T1*). Data are means ± SEM. ***: p<0.001; **: p<0.01*; p<0.05.

## DISCUSSION

Chromosomal translocations are the result of two simultaneous DNA double-strand breaks in different chromosomes followed by reciprocal erroneous rejoining of DNA ends. In the case of balanced translocations, frequent in lymphoid malignancies, two aberrant chromosomes are formed and called derivative (der) chromosomes (Lieber, 2016). Recurrent chromosomal translocations are found in blood and certain solid tumors.

After a translocation one of the most recurrent effects is the overexpression of an oncogene following its juxtaposition with a highly expressed gene. This is the case for example for Burkitt lymphoma, Follicular lymphoma or Mantle cell lymphoma where *MYC*, *BCL2* and *CCND1*, respectively, result overexpressed following their erroneous joining next to an immunoglobulin gene locus. These recurrent chromosomal aberrations often serve as a marker of the tumor.

The presence of a translocation does not inevitably mean oncogenic transformation, indeed genomic translocations are also found in healthy individuals, thus meaning that additional mutations and alterations are required to produce a malignant phenotype (Aplan, 2006). At the same way also oncogene overexpression occurring after the translocation is not enough to produce a malignant phenotype and additional alterations should arise (Grumolato and Aaronson, 2014).

This is true for MCL for which it has been demonstrated that *CCND1* (located on the chromosome 11) alone is not enough to induce the malignant transformation and additional alterations involving cell cycle, DNA damage response or epigenetic modifications are required (Peterson et al., 2019). An understanding of the events and the mechanisms leading to lymphomagenesis is still lacking.

We have found that common genes are deregulated *in vitro* and in MCL patients on the chromosome 11 and half of them was concentrated in a relatively small region of 15 Mb next to the translocation breakpoint.

Among the deregulated genes we found genes implicated in DNA damage and repair (*ATM*, *MRE11* and *RAD9A*) or others that has already been described as deregulated or markers or mutated in lymphomas or other cancers (*AHNAK*, *CD6*, *CTSF*, *CPT1A*, *MEN1*) or associated to B-cells proliferation (*CYB561A3*, *MAP3K11*) (Hartmann et al., 2008; Hoster et al., 2008; Knackmuss et al., 2016; Melone et al., 2018; Pasqualucci et al., 2011; Rizzatti et al., 2005; Thieblemont et al., 1999).

Consistently with other studies, also the glutathione S-transferases P1 gene (*GSTP1*) gene was upregulated. Proteins from GSTs family play an important role in the cell detoxification from xenobiotics that may be carcinogenic (Seidegård and Ekström, 1997) and *GSTP1* expression level deregulation has already been associated to prostate, breast, liver, renal, and endometrial carcinomas (Yuan et al., 2008) and to MCLs (Bennaceur-Griscelli et al., 2004; Bosch et al., 1994).

According with other findings, also genes located on chromosomes other than the 11 were found deregulated as *TP53*, *SOX11*, *BCL* family members and *RB1* (Kienle et al., 2007; Meggendorfer et al., 2013; Rummel et al., 2004; Sakhdari et al., 2019)

The obtained results were also confirmed by data mining of transcriptomic data from Gene Expression Omnibus (GEO) of 15 conventional MCL patients' samples composed of Peripheral blood CD19+ tumor cells (GSE16455) and of 3 naive B-cells samples (GSE12366). We have found that the expression of genes located in two short regions (4 Mb) on chromosomes 7 and 9 and one long region (15 Mb) on the chromosome 11 in proximity of the translocation breakpoint, mainly located on the der11 were significantly deregulated. This event was specifically occurring in MCL after the t(11;14), indeed performing the same analysis on data obtained from Follicular Lymphoma cells, the expression of the genes located in the same regions was not significantly changed.

For *CCND1* overexpression mechanisms in MCL, different hypothesis have been formulated (Allinne et al., 2014; Liu et al., 2008). It is assumed that it depends on the potent Eµ enhancer located upstream of *CCND1* in the *IGH* locus even if the distance between the two partners could be several hundred kilobases (Degan et al., 2002; Greisman et al., 2012). Our laboratory previously proposed that the *CCND1* transcriptional activation in MCL was related to the repositioning of the rearranged *IGH*-*CCND1*-carrying chromosome (der14) in close proximity with the nucleolus, a nuclear territory with abundant nucleolin acting as transcriptional factor and PolII molecules.

Translocations can lead to large scale modification of chromosome territory position affecting the expression of many different genes (Harewood et al., 2010).

Analysis of the nuclear positioning and transcriptional modifications of the der11, one of the chromosomes generated by the translocation t(11;14), carrying a large portion of the chromosome 11 (~71Mb) and a small part of chromosome 14 (~10 Mb), revealed that surprisingly the presence on this chromosome of a small fragment of the chromosome 14 is enough to induce a large movement of a large portion of the chromosome 11 in the nuclear space. We observed that the der11 relocates in close proximity to the nucleolus where the nucleolin, component of the LR1 transcription factor, can directly bind chromatin and lead to the deregulation of gene expression. We found the same transcriptional deregulation and nuclear remodelling in four MCL patients in leukemic phase.

In summary, in the present and previous works from our laboratory, we provided novel proofs of genes as possible markers for MCL other than *CCND1* and describe large scale chromatin modification leading to a nucleolin dependent massive gene deregulation in specific regions of der14 and der11 chromosomes.

## ACKNOWLEDGEMENTS

This research was supported by grants from the ANRS, INSERM (ENVIBURKITT) and La Ligue Contre le Cancer (M27231). We gratefully acknowledge the imaging and cytometry platform of the Gustave Roussy Institute for the technical support.

## MATERIALS AND METHODS

### Cell cultures

The human Mantle cell lymphoma (MCL) cell line GRANTA-519 waskept in culture in RPMI 1640 medium supplemented with 10% heat-inactivated FBS 2 mM L-glutamine and 1% penicillin/streptomycin antibiotics (Gibco®, Thermo Fisher Scientific, Carlsbad, CA, USA). For cultivating the lymphoblastoid cell line (LCL) RPMI8866 this medium was further supplemented with 2% Glucose, and 1mM Sodium pyruvate (Gibco®, Thermo Fisher Scientific, Carlsbad, CA, USA).

Peripheral blood mononuclear cells (PBMCs) were isolated from blood of MCL patients by Pancoll (PAN biotech, Aidenbach, Germany, lymphocytes separation medium) density gradient centrifugation. B lymphocytes were obtained by negative cell selection using the MagniSort Human B-cell enrichment kit II (Thermo Fisher Scientific, Carlsbad, CA, USA) according to the manufacturer’s protocol. They were kept in culture for 24h into RPMI medium supplemented with 10% heat-inactivated FBS and1% penicillin/streptomycin antibiotics(Gibco®, Thermo Scientific, Carlsbad, CA, USA)before collecting them for further analysis.

Human Embryonic Kidney (HEK-293) cells were cultivated in DMEM (Gibco®, Thermo Scientific, Carlsbad, CA, USA) supplemented with 10% FBS,2mM glutamine and 1% penicillin/streptomycin.

All cells were cultured at 37°C in a humidified 5% CO2 atmosphere.

### Mantle cell lymphoma patients

Whole blood samples were obtained from four individuals followed in the Department of Medical Oncology, Institut Gustave Roussy, Villejuif, France. Blood samples were collected in agreement with French law and informed consent was obtained from all subjects.

### RNA isolation, library preparation and RNA sequencing

RNA was extracted from LCL RPMI8866, GRANTA-519 and B-cells purified from four MCL patients using the NucleoSpin® RNA II kit according to the manufacturer’s protocol (Macherey-Nagel, Oensingen, Switzerland). A minimum of 2µg per samples were used for analysis. Samples were processed and analyzed at NovoGene Co., Beijing, China. Quality and quantity were assessed with Qubit Fluorometer, poly-A enriched library prepared, samples sequenced using an Illumina PE150 and analyzed.

### Replication timing analysis

LCL RPMI8866 and GRANTA-519 were pulse labeled with BrdU and sorted into two S-Phase fractions (early and late S-Phase). BrdU labeled nascent DNA from early and late fractions were immunoprecipitated with an anti-BrdU antibody, amplified, differentially labeled, and co-hybridized onto a whole human genome oligonucleotide microarray. The log_2_-ratio of the abundance of each genomic probe in the early and late S-phase fractions generates a replication-timing profile that reveals early and late replicated domains.

Replication timing analysis was carried out as previously described (Hassan-Zadeh et al., 2012). Briefly, immunoprecipitated nascent strands were amplified by whole-genome amplification kit (Sigma Aldrich, St Louis, MO, USA). After amplification, early and late nascent strands were labeled with Cy3 and Cy5 ULS molecules using the Genomic DNA labeling Kit (Agilent, Santa Clara, CA, USA) following manufacturer's instructions. The hybridization was performed according to the manufacturer instructions on human CGH microarrays that cover the whole genome. Microarrays were scanned with an Agilent's High-Resolution C Scanner using a resolution of 2 µm and the autofocus option. Feature extraction was done with Feature Extraction 9.1 software (Agilent, Santa Clara, CA, USA). For each experiment, the raw data sets were automatically normalized by the Feature extraction software. Analysis was performed with Agilent Genomic Workbench 5.0 software. The log2-ratio timing profiles were smoothed using the Moving Average option of the Agilent Genomic Workbench 5.0 software with the linear algorithm and 200 kb windows.

### Immuno-3D-FISH

“RainbowFISH” probes (Empire Genomics, Buffalo, NY, USA)) used for hybridization and staining of human chromosome loci 14q32.33 (*IGH* gene locus) and 11q13.2 (*GSTP1* gene locus) were labeled with SpectrumOrange and SpectrumGreen respectively. Cells were immunofluorescently stained with the rabbit anti-nucleolin antibody (Sigma-Aldrich). Nuclei were stained with DAPI diluted into the Vectashield mounting medium (Vector Laboratories, Burlingame, CA, USA). Immuno-3D-FISH was performed as previously described (Germini et al., 2017).

### Microscope image acquisition and analysis

Images were acquired using a TCS SP8 confocal microscope (Leica Microsystems, Berlin, Germany) with a 63X oil immersion objective. Z-stacks were acquired using a frame size of 1024×1024, and 0.5 µm z-steps, with sequential multitrack scanning using the 405, 488, 543, 633 nm laser wavelengths for detecting DAPI (cell nuclei), SpectrumGreen (*GSTP1*), SpectrumOrange (*IGH*) and far-red (Nucleolin). Radial position and distance from the closest nucleolus of gene loci was calculated using the Bitplane® Imaris (Zurich, Switzerland) program.

### HEK cells transfection and sorting

400,000 cells were plated in 6-well plate in complete DMEM medium. The day after when cells reached approximately 70% of confluence they were transfected with 3µg of the PEGFPC1 plasmid (Addgene #28176) using the Viafect reagent (Promega, Madison, WI, USA) following manufacturer's protocol. Transfection was carried out for 48h and then GFP+ cells sorted using the ARIA III cell sorter (Becton Dickinson, Franklin Lakes, NJ, USA).

### RT-qPCR

Total RNA was extracted from cells using the NucleoSpin® RNA II kit according to the manufacturer's protocol (Macherey-Nagel). Sixty nanograms of total RNA were reverse transcribed by using the RevertAid H Minus First Strands cDNA Synthesis Kit with oligo(dT) primers (Thermo Scientific). The obtained cDNA was amplified using specific primers (**Supplementary Table 1**) and the PowerUp SYBR Green Master Mix (Thermo Scientific). Expression of target genes was analyzed using the 2^-ΔΔCt^ method with normalization with *GAPDH* and comparison of expression between non transfected (Control) and GFP+ cells. For quantification, expression levels were set to 1 in control.

### Chromatin Immunoprecipitation (ChIP)-seq

Thirty millions cells were fixed with 1% formamide for 1 minute in a rotating wheel. Fixation reaction was stopped by adding 0.125M glycin. Cells were then pelleted at 4°, 1400xg for 10 minutes and washed 3 times in ice cold 1XPBS. At the third wash, 0.5%PIC and 0.5mM of PMSF were added. Cells were pelleted and resuspended in 1 mL of lysis buffer (10mM Tris-HCl, 150mM NaCl, 0.5mM EDTA, 0.5% NP-40, 0.5% PIC, 0.5mM PMSF, water). Cell lysate was transferred to a dounce homogenizer to allow the separation between cell cytoplasm and nuclei and dounced 10 times. Nuclei are then pelleted by centrifugation at 5000xg for 5 minutes at 4° and resuspended in nuclear lysis buffer (50mM Tris-HCl, 10mM EDTA, 1% SDS, 1% PIC, 1mM PMSF, water). Nuclei were sonicated in ice in a Vibra Cell sonicator (Newtown, CT, USA) with25 cycles of 30 sec pulse-on, 30 sec pulse-off, 75% amplitude. The non-solubilized material was removed by centrifugation at 15,000g for 10 min. The desired size of chromatin fragments (100-500 bp) was monitored by electrophoresis in a 1% agarose gel after reverse crosslinking (dilution 1:4 in water and adding 0.250M of NaCl keeping at 65° overnight in a thermalcycler), treatment with 100µg/ml RNase A (37° for 15 minutes) and 50µg/ml proteinase K and purification using the PCR cleanup kit (Macherey-Nagel).

Chromatin immunoprecipitation was performed as following: 25 µg of chromatin solution was incubated at 4° overnight with 25µl of the PrG-Dynabeads (Thermo Scientific),3µg of antibody (rabbit-anti Nucleolin, Sigma-Aldrich), ChIP Buffer I (Active Motif, La Hulpe, Belgium), 1X PIC and water. The following day the beads were washed once with ChIP Buffer I and twice with ChIP Buffer II (Active Motif) before elution with 50mM Glycine (pH 2.8) for 15 minutes at room temperature. Elution was stopped by adding Tris base 1M (pH10.4). By using a magnet, supernatant was kept, reverse cross-linked, purified and quantified by using a Nanodrop (Thermo Scientific).

Before sequencing, effective ChIP was checked via PCR amplification of known positive and negative targets of the nucleolin by normalizing the results using the input sample.

A minimum of 100ng of ChIP DNA were sent to NovoGene for sequencing where quality and quantity were assessed with Qubit Fluorometer, library prepared, samples sequenced using an Illumina PE150 and analyzed.

### Statistical analyses

The Student t-test was used to compare means. All tests were performed using the GraphPad Prism 5 software (GraphPad software Inc., La Jolla, CA, USA).

