## Supplementary figures and images for "Large-scale nuclear remodeling and transcriptional deregulation occur on both derivative chromosomes after Mantle Cell Lymphoma chromosomal translocation"

### Supplementary Figures

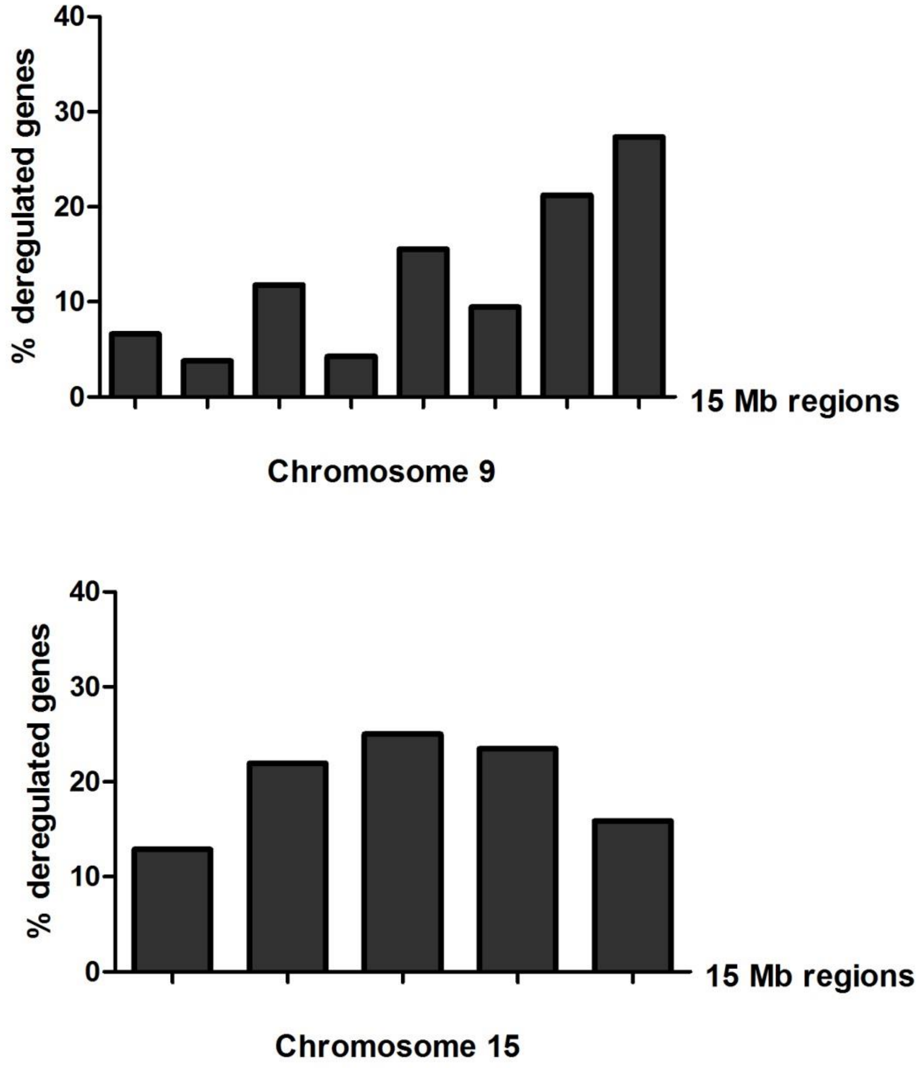


Supplementary Figure 1

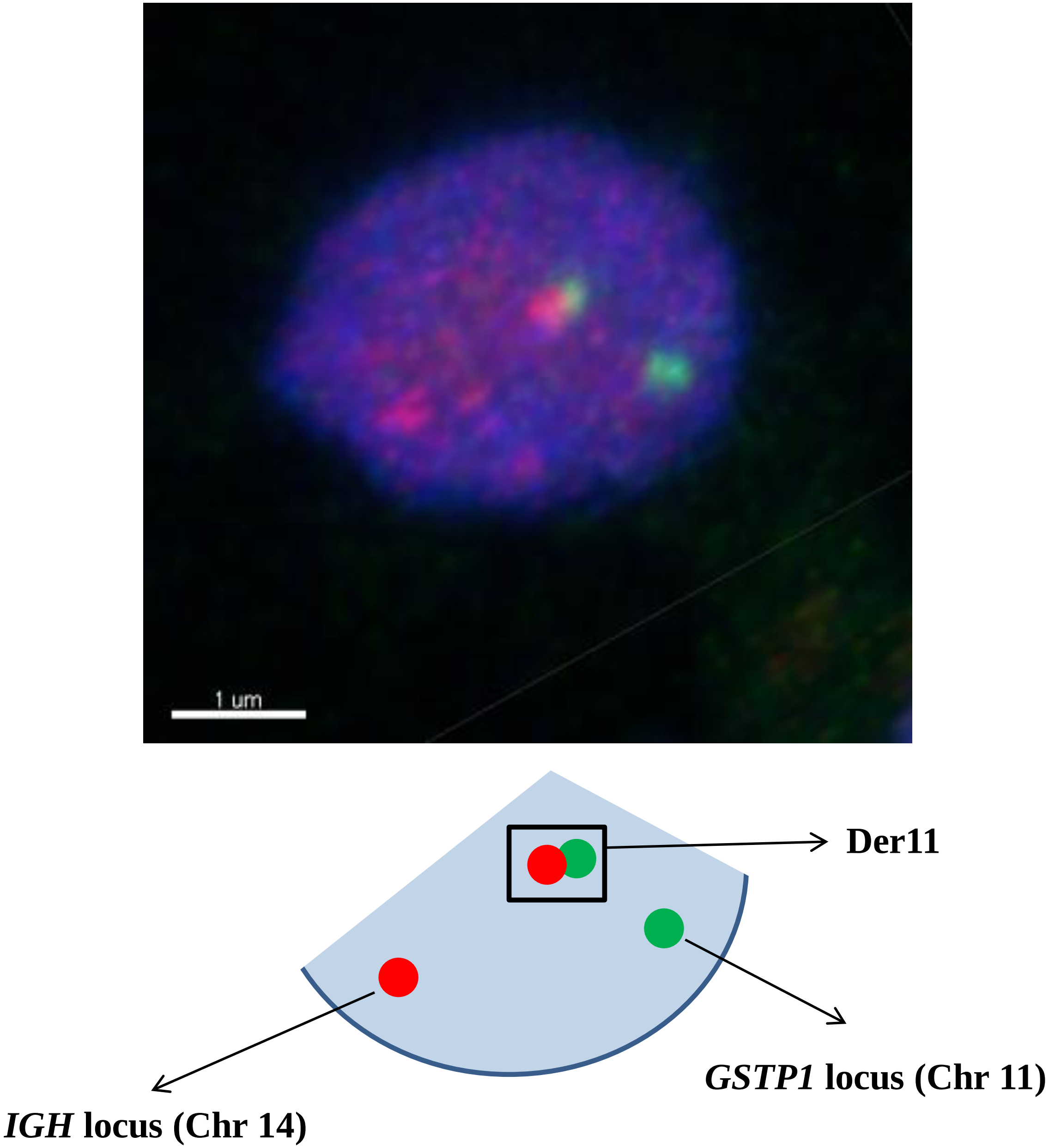


Supplementary Figure 2

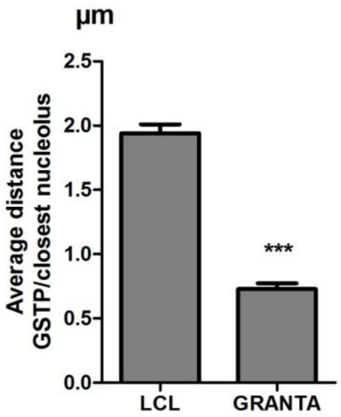


Supplementary Figure 3

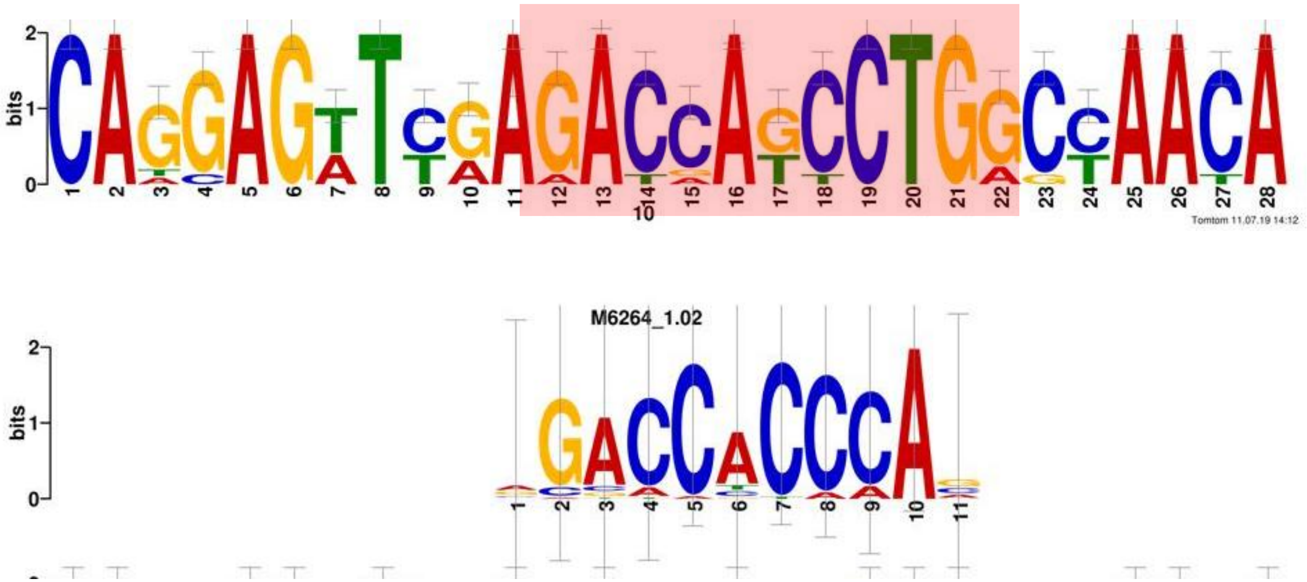


Supplementary Figure 4
