## Supplementary Table 1 for "Large-scale nuclear remodeling and transcriptional deregulation occur on both derivative chromosomes after Mantle Cell Lymphoma chromosomal translocation"

**Supplementary Table 1.** Primer used for qRT-PCR

| GENE NAME | PRIMER F 3'-5' | PRIMER R 3'-5' |
|---|---|---|
| CD6 | GTGACAGTGAGTGGTAC | CCATTACGGGAGGGCATAT |
| AHNAK | GTGACCGAGATTCCCGACGA | GAGGAGACAACCCGGGAGCT |
| RASGRP2 | GCATGTTCCTCATGATGCAC | GAGGAGACAACCCGGGAGCT |
| PYGM | GCATCAAAAGGGAGCCCAAT | GCCTTGCCTCCAATCATGA |
| CTSF | GACTGTGACAAGATGGACAA | CCACGGAGTCATTGATGTAGA |
| CPT1A | TCCAGTTGGCTTATCGTGGTG | CTAACGAGGGGTCGATCTTGG |
| DAK | TTGTCACTTCCTTCTGCCTT | ATGCCTTATCTCTGAAGCCC |
| RTN3 | TCTGTCCAGTTTTCAGCACT | ATTGTGGGGCAGGAAGATAG |
| SOD1 | CCTCTATCCAGAAAACACGG | CCAAACGACTTCCAGCG |
| RUNX1T1 | AACCGATGAGGTTCTCCTGA | CGGCTTTAAAATGGGGTCTT |
| NUCLEOLIN | GATCACCTAATGCCAGA | CAAAGCCGCCTCTGCCTCCACCAC |
| GAPDH | CTGCACCACCAACTGCTTAG | AGGTCCACCACTGACACGTT |
