## Supplementary figure Legends for "Large-scale nuclear remodeling and transcriptional deregulation occur on both derivative chromosomes after Mantle Cell Lymphoma chromosomal translocation"

**Supplementary Figure 1.** Percentage of deregulated genes in non-overlapping 15mb regions of the chromosome 9 and 15. The percentage was calculated on the total number of upregulated genes of the chromosomes 9 and 15.

**Supplementary Figure 2.** 3D-FISH image representing the typical GRANTA-519 cell. It displays *IGH* loci in red and *GSTP1* loci in green. Colocalization between one allele of each gene represents the Der11 chromosome (as shown in the schematic representation below the 3D-FISH image). One non-colocalized allele of each locus is also present. The 3D-FISH image is obtained by the Imaris Software.

**Supplementary Figure 3.** Average distance from the nucleolus of the *GSTP1* loci on the der11 chromosome (“GRANTA”) and on the normal Chromosome 11(“LCL”). A minimum of 216 loci were analyzed.

**Supplementary Figure 4.** Motif discovery from Nucleolin peaks obtained in ChIP-seq analysis. Identification of statistically significant represented sequences present in ChIP-seq Nucleolin binding loci using the motif discovery tool MEME. The consensus sequence previously reported is highlighted in red (upper panel) and the novel one is shown in the bottom panel.
